# Biofilm formation on agricultural waste pretreated with cold low-pressure nitrogen plasma and corona plasma discharges

**DOI:** 10.1101/299172

**Authors:** Ravit Farber, Inbal Dabush-Busheri, Gilad Chaniel, Shmuel Rozenfeld, Edward Bormashenko, Victor Multanen, Rivka Cahan

## Abstract

Agricultural waste (AW) was pretreated with cold low-pressure nitrogen plasma (LPD) and corona atmospheric plasma discharges (CAPD), in an attempt to increase the bacterial attachment and biofilm formation. Biofilm formation was examined in the presence of exogenously added *P. putida* and *B. cereus* as well as in a sterile medium where only the indigenous bacteria which grow naturally on the wood surface could form biofilm. The exposure of AW to (LPD) led to a 3.5-fold increase in biofilm formation of the exogenously add *P. putida* F1 in MMT (minimal medium supplied with toluene) and a 1.6-fold increase in MMG (minimal medium supplied with glucose) compared to the untreated AW. The increase in biofilm formation was also observed with the exogenously added *B. cereus* or with indigenous bacteria that grow naturally on the AW. The effect of the CAPD on biofilm formation was weak. SEM analysis of the LPD-treated AW showed an increase in surface roughness, which we assume is one of the reasons for the enhancement of the biofilm formation. The apparent contact angle of a sessile drop on the surface of LPD-treated AW as well as on the bacterial layer showed their hydrophilic nature. In conclusion, the increase in biofilm formation of the exogenously added *P. putida* or *B. cereus* was due to the LPD treatment.

**Importance:** To the best of our knowledge, this is the first study to describe the effect of wood plasma treatment on biofilm formation. This technology can be further implemented for bioremediation of contaminated soils.

## Introduction

Plasma is one of the four forms of the naturally occurring matter states, and is mostly comprised of ions and electrons (1). Plasmas are considered as an ionized gas which can be divided into two types according to their gas temperature, “Hot” plasmas (near-equilibrium plasmas) and “Cold” plasmas (non-equilibrium plasmas). Hot plasmas are characterized by very high temperatures of electrons and heavy particles, whereas cold plasmas are composed of low temperature particles and relatively high temperature electrons. Hot plasmas include electrical arcs, plasma jets of rocket engines, thermonuclear reaction generated plasmas, etc. Cold plasmas include low pressure direct current (DC), radio frequency (RF) discharges (silent discharges), and discharges appearing in fluorescent (neon) illuminating tubes. Corona plasma discharges are also identified as cold (however, atmospheric pressure) plasmas (2).

Cold plasma discharges are effectively used for a plethora of technological processes, including modification of chemical and physical properties of organic (synthetic and natural) surfaces (3), polymerization (4), antibacterial treatment of non-woven fabrics (5) and advanced agriculture technologies (6), etc.

Due to the relatively small energy of particles, inherent to the cold plasma discharges, the depth of influence (penetration) of these discharges into organic substrates is nano-scaled (7).

It is generally accepted that plasma treatment creates a complex mixture of surface functionalities which influence physical and chemical properties of a surface, usually resulting in its hydrophilization (8). The plasma inspired changes in physical and chemical properties of surfaces are often (but not always) lost with time (9). This process is called “hydrophobic recovery” and is much more pronounced for surfaces of synthetic polymers than for biological surfaces (6, 9).

Cold plasma utilization was examined for bacterial eradication (10), for elimination of soil bacteria (11) and for increasing seed germination and crop yield (12). Several studies reported that wood plasma treatment led to improved adhesion properties and wettability to liquid (13), which are important for the contact of adhesives (14) to wood composites (15).

Biofilms are bacterial communities attached to surfaces. Within the biofilm there are quorum sensing signals leading to up and down regulation of gene expression that enables adaptation to various environmental organic pollutants (16). Biofilms have the ability to survive in nutrient deficient conditions (17) are highly resistant to hydration (18) and have mechanical stability (19). Immobilized bacteria for bioremediation is a cost-effective and attractive method for degradation of various hazardous environmental pollutants (20). These advantages have encouraged researchers to developed methods for bacterial adhesion to surfaces, which is the first and critical step for biofilm formation. Factors influencing bacterial adhesion to substrata are the substratum topography and wettability as well as bacterial cell surface charges and structures such as flagella, capsules, pili and fimbriae (21, 22). Substratum roughness, such as pores and scratches, increase the surface area for bacterial adhesion (23). The degree of surface wettability describes the contact ability of a liquid to a solid substratum which is influenced by the interaction between the fluid and the solid substratum (24). Successful adherence of bacteria to a substratum occurs when dipole, ionic, hydrogen, electrostatic, hydrophilic or hydrophobic interactions between the bacterial surface and the substratum are strong enough that less than 5 nm separate between the bacterial cell and the substratum (21). Modification of the substratum surface is an important way for increasing bacterial attachment and biofilm formation for bioremediation process. The modification methods should be scalable, economical and environmentally friendly (25).

Proposed surfaces for biofilm formation includes: polymer membranes (26), activated carbon (27), glass surfaces (28), coal bottom ash, geo-textile sheets and polyethylene terephthalate fibers (29, 30).

In this study, the substrata for biofilm formation were thin branches of a local pine tree, which are considered as agricultural waste (AW). The AW was pretreated by cold low-pressure nitrogen plasma and corona discharges in an attempt to increase bacterial attachment and biofilm formation. Biofilm formation was examined in the presence of exogenously added *P. putida* and *B. cereus* as well as in a sterile medium where only the indigenous bacteria, which grow naturally on the wood surface, could form the biofilm. To the best of our knowledge, this is the first study to describe the effect of wood plasma treatment on biofilm formation. This technology can be further tested for increasing bioremediation of contaminated soil.

## Materials and Methods

### Agricultural waste

Thin branches of a local pine tree served as the agricultural waste. The thin branches were cut to pieces of 0.06 g. For the biofilm experiments, three pieces were added to each 50 mL tube.

### Biofilm growth conditions

*P. putida* F1, strain 6899 (DMSZ) was grown in mineral medium (MM) (30) supplied with 300 mg L^-1^ toluene (MMT) or in MM supplied with 1% glucose (MMG) in sealed bottles to the log phase at 30°C with shaking at 170 rpm. The culture optical density was measured using a GENESYS 10S UV-Visible spectrophotometer (Thermo Scientific, USA) at 590 nm. For biofilm formation on the AW, the cultures were diluted in MMT or MMG to 0.15 OD in 50 mL tubes containing three pieces of AW, and incubated at 30°C, in a static mode for 48 h.

### Biofilm viability measurements

The biofilm on the AW was washed three times with phosphate buffered saline (PBS) to remove the planktonic bacterial cells. The biofilm viability was measured by adding 2 mL of 3-(4,5-dimethylthiazol-2-yl)-2,5-diphenyltetrazolium bromide (MTT) solution (5 mg mL^-1^ in 0.1 M PBS, pH 7.4) for 3 h at room temperature in the dark (29). The reduced MTT salts was measured at 540 nm using a spectrophotometer. When the absorbance was higher than 1 OD, the sample was diluted and reexamined.

### Plasma treatment

#### Cold low pressure nitrogen plasma (Low pressure discharge)

The AW was exposed to a radiofrequency RF (13.56 MHz) inductive pressure nitrogen plasma discharge (created by the Harrick Plasma cleaner/sterilizer PDC-3XG (MMT Corp. USA). The low-pressure discharge (abbreviated further LPD) is sustained by an RF oscillating electric field, generated in the gas region at a pressure (P) of *ca* 1.0 Torr and a power of 18 W for 1 min (Fig. 1). Nitrogen 99.999% was supplied by Oxygen and Argon Works Ltd, Israel. Immediately after the cold nitrogen plasma treatment, the AW was placed in distilled water to preserve its surface activation (31).

**Figure 1:**
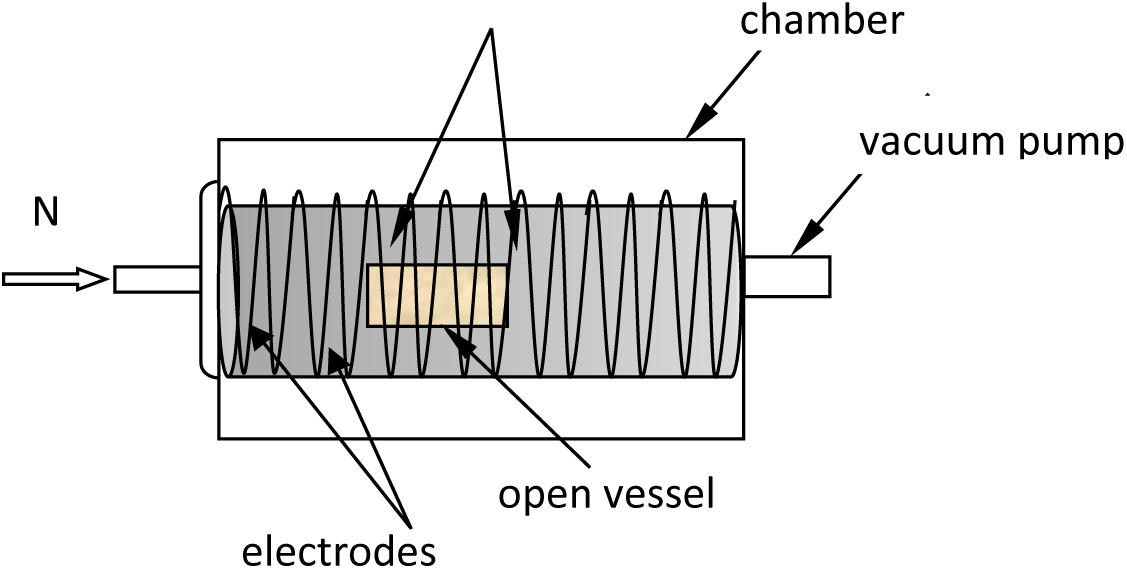
Sketch of the N_2_ plasma treatment of the agricultural wastes is depicted.

#### Corona plasma discharge (atmospheric pressure discharge)

Plasma installation, i.e. MultiDyne 1000 unit (supplied by 3DT Co. USA) generated an air atmospheric pressure corona plasma discharge (abbreviated further CAPD) at ambient conditions. The corona plasma discharge was created by applying high voltage (about 12 KV•cm^-1^) pulses at a freguency of 50 Hz in the clearance of *ca* 5 mm between electrodes. The corona plasma discharge was transferred with a stream of air to the treated surface. The air stream was created by the fan (supplied by 3DT Co. USA).

### Bacterial cell surface and agricultural waste hydrophobicity

#### Preparation of the bacterial cell for measurement of the apparent contact angle of water on filtered bacterial layer

*P.putida* F1 as well as *Bacillus cereus* were cultivated in MMT or MMG at the end of the log phase. The cultures were centrifuged twice in water and filtered on 0.45µm membrane (Sartorius). The layer of *P. putida* F1 on the filter membrane was dried in a desiccator for 1 h (32).

#### Preparation of the bacterial cell for measurement of the apparent contact angle of water on highly concentrated bacterial layer placed on glass slide

The cultures (50 mL) were grown on MMT or MMG to the log phase and washed as described for the Van Loosdrecht method (32). The bacterial sediment was suspended in 200-300 µL water and placed on a glass slide to a homogenous thick suspension, followed by incubation at 30°C for 2 h.

The apparent contact angle of a sessile drop (1 µL) of distilled water on the filtered bacterial layer as well as on the highly concentrated bacterial layer placed on the glass slide was examined using a Ramé-Hart Advanced Goniometer Model 500 (6).

#### Measurement of the apparent contact angle of water on the agricultural waste surface

The apparent contact angle of water on the AW before and after the LPD treatment, was measured with the Ramé-Hart Advanced Goniometer Model 500 (6).

#### Scanning electron microscopy (SEM) analysis of the agricultural wastes

The sample was mounted onto an aluminum stub with a two-sided adhesive carbon tape followed by coating the sample with a thin (20 nm) gold layer using QUORUM Technologies Q150 T ES. Scanning was performed with an ultra-high resolution SEM

##### Statistic

Data are expressed as means ± SD (standard deviation) of two different experiments and three replicates. The results were statistically analyzed using one-way analysis of variance (ANOVA). Values were considered significant at P < 0.05.

## Results and Discussion

### Biofilm formation on agricultural waste pretreated with LPD

LPD-treated and untreated agricultural wastes (AW) were added to a culture of *P. putida* F1 (0.15 OD) grown in MM supplied with toluene. The controls were LPD-treated and untreated AW that were added to a sterile MMT (without exogenous *P. putida* F1) and AW that was treated only at 1 Torr (vacuum treatment) without plasma treatment. After 48 h, the AW was washed and the biofilm viability was measured by the calorimetric analysis, MTT (Fig 2 A). In this assay the hydrogenases of the exogenous *P. putida* F1 and the indigenous bacteria, which grow naturally on the surface of the AW, reduce the MTT tetrazolium salt reagent to a purple solution that can be measured spectrophotometrically. The results show that the biofilm viability on untreated and LPD-pretreated AW in the presence of the exogenous bacteria *P. putida* F1 were 0.55±0.11 and 1.92±0.09 OD 540 nm, respectively. When the untreated and the LPD-pretreated AW were added to a sterile MM in which only the indigenous bacteria which grow naturally on the AW can formed a biofilm, the biofilm viability was 0.36±0.02 and 1.54±0.20 OD at 540 nm, respectively. In the presence of the exogenous *P.putida*, biofilm viability on the AW that was treated only by the vacuum was 0.52±0.09 OD, which is similar to the biofilm viability on the untreated AW (0.55±0.11 OD 540 nm).

**Figure 2:**
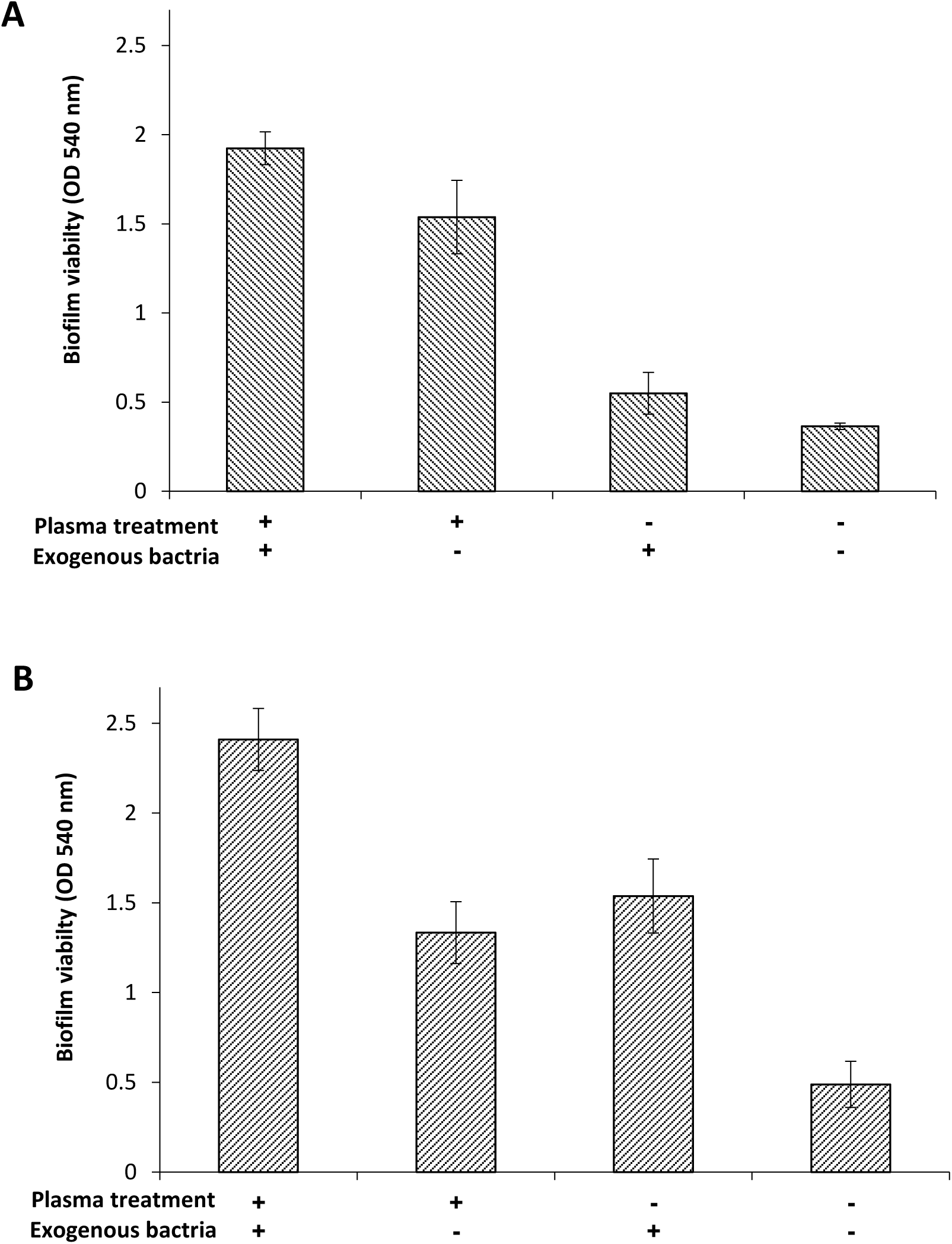
Biofilm viability on agricultural waste in the presence of toluene A or glucose B. Plasma pre-treated agricultural waste (+); untreated agricultural waste(-). In the presence of exogenous bacteria, *P. putida* F1 (+) or without (-).The P-value between the biofilm of the exogenous bacteria on the treated and the untreated AW is P < 0.05. Same P-value for the biofilm of the indigenous bacteria.

The same experiment was performed, but in the presence of glucose (Fig 2 B) instead of toluene. In the presence of the exogenous bacteria the biofilm viability on the LPD-pretreated AW was (2.41±0.17 OD 540 nm), which is 1.6-fold higher than the biofilm viability on the untreated AW. The plasma treatment also influenced the biofilm formation of the indigenous bacteria. Biofilm viability on the LPD-treated AW (1.54±0.20 OD) was 4.2-fold higher than on the untreated AW.

In conclusion, the LPD-treatment significantly increased the biofilm formation of the exogenous and indigenous bacteria when grown on toluene or glucose as the sole carbon source.

Biofilm formation was also examined on AW pre-treated with LPD that was added to a culture of *B. cereus* as described for *P. putida*. The results presented in Figure 3 A and B, indicate that the biofilm viability on the LPD-pretreated AW in the presence exogenous *B. cereus* grown on toluene as well as on glucose was 0.96±0.04 and 2.06±0.27 OD 540 nm, respectively, which is 1.3-fold higher than the biofilm viability on the untreated AW. The biofilm viability of the indigenous bacteria on the plasma pretreated AW was 1.3 fold higher on the untreated AW (Fig 3 B).

**Figure 3:**
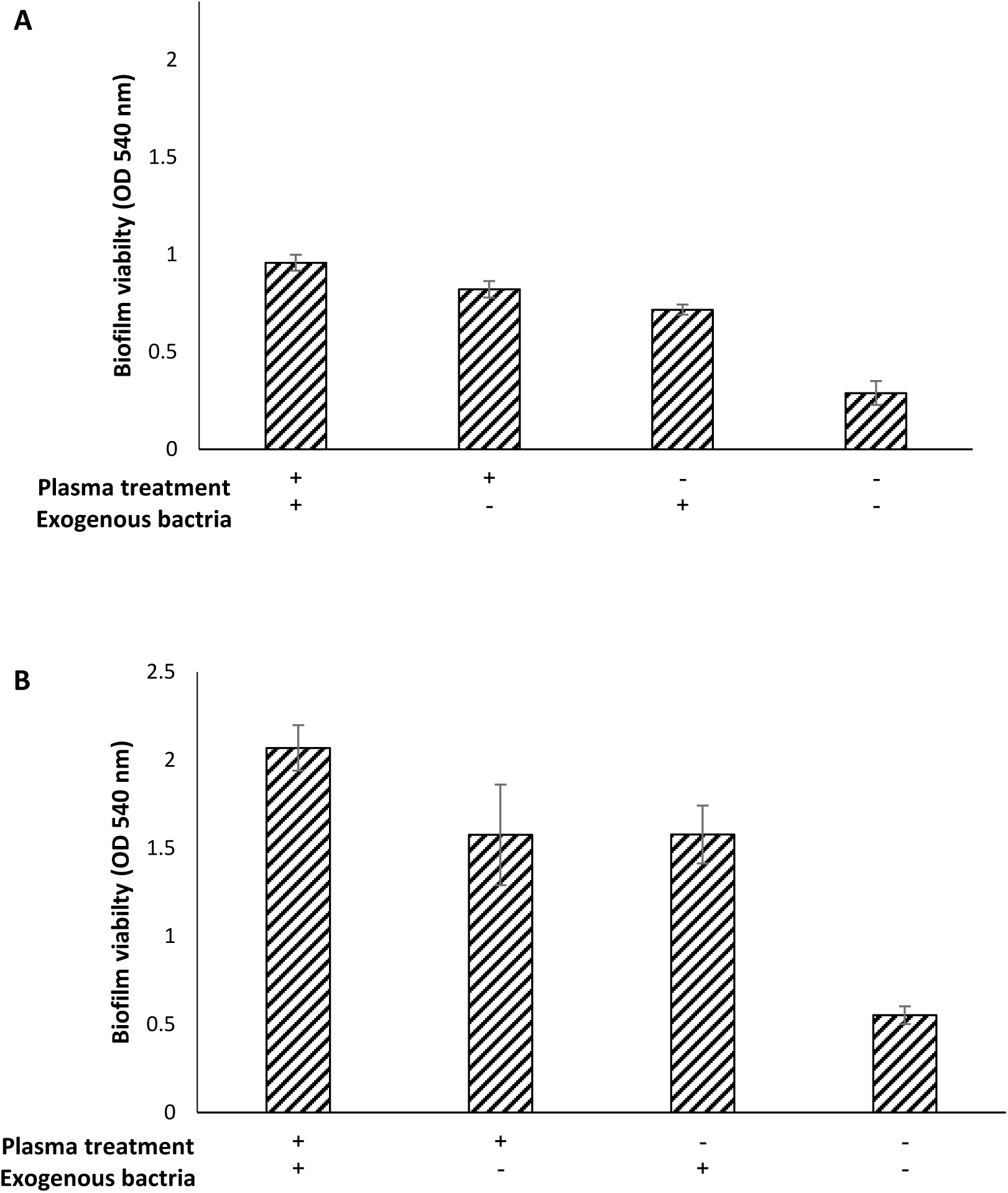
Biofilm viability on agricultural waste in the presence of toluene A or glucose B. Plasma pre-treated agricultural waste (+); untreated agricultural waste(-). In the presence of exogenous bacteria, *B. cereus* (+) or without (-).The P-value between the biofilm of the exogenous bacteria on the treated and the untreated AW is P < 0.05. The same P-value was obtained for the biofilm of the indigenous bacteria.

### Biofilm formation on corona atmospheric plasma discharge pre-treated agricultural waste

AW was treated by corona atmospheric plasma discharge (CAPD), for 1 min at distances of 2 and 5 cm and placed in tubes containing MMT with and without *P. putida*. After 48 h the agricultural waste was washed and the biofilm viability was measured by MTT analysis (Fig 4). It can be seen that in the presence of *P. putida* the biofilm viability on AW that was treated using CAPD at a distance of 2 and 5 cm, was 0.33±0.03 and 0.79±0.02 OD 540 nm, respectively. In the absence of exogenous bacteria, the biofilm viability was only 0.15±0.01 and 0.65±0.05 OD, respectively. Since the effectiveness of the LPD on biofilm formation was markedly higher than that of the CAPD (explanation of this phenomenon will be discussed later), further experiments of biofilm formation were performed only using the LPD treatment.

**Figure 4:**
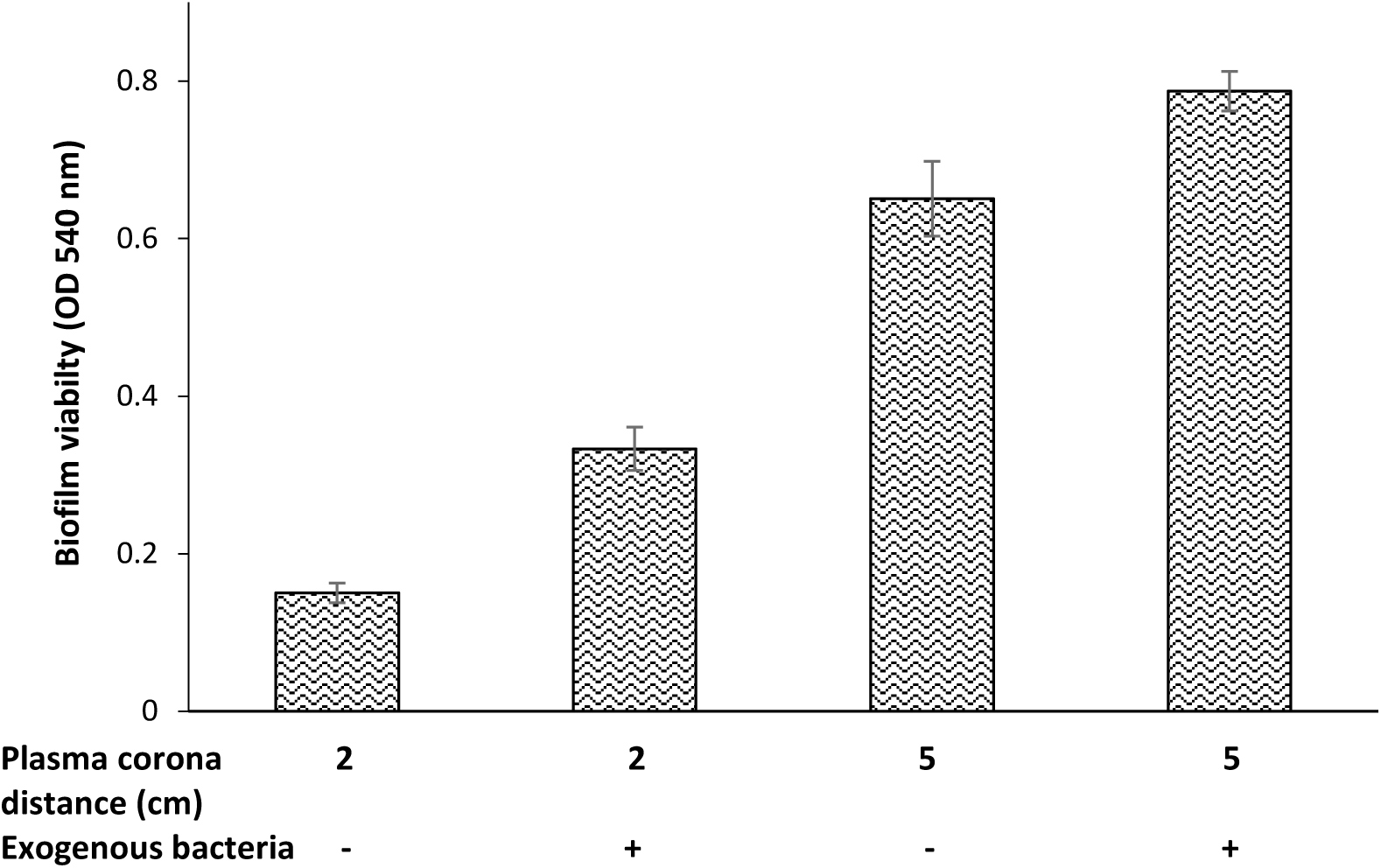
Biofilm viability on AW pre-treated with corona plasma atmospheric discharge at a distance of 2 and 5 cm; when the agricultural waste was placed in tubes containing a culture of *P. putida* F1 (+) or without *P. putida* (-).

### Morphology of the xylem vessels after plasma treatment

The AW was treated using LPD as described in the Materials and Methods, and was immediately covered by a thin layer of gold and analysed using SEM. The control for the LPD was a sample of AW that was treated only by vacuum without exposure to plasma. In addition, a sample of AW was treated using CAPD at a distance of 2 cm in order to elucidate the reason of the lower effectiveness of CAPD on biofilm formation. SEM analysis was performed on the internal and external surface of the AW (Fig 5). The internal untreated AW xylem vessels are organized in straight lines and the average vessel width is 10.4±1.4 µm (Fig 5 A). Irregularity of the xylem vessels with the increase in the surface area was found in the AW treated using LPD. The width of the xylem vessels showed a high diversity of between 4.4 to 26.4 µm, with a mean width of 7.6±5.7 µm (Fig 5 B). The image of the AW that was treated only by a vacuum shows no significant influence on the xylem vessels’ topography compared to the untreated AW, with a mean xylem vessel width of 13±3.9 µm (Fig 5 C). However, when the agricultural waste was treated using CAPD the width of the xylem vessels increased to 21.3±4.3 µm and the surface of xylem vessels edge became smooth (Fig 5 D). The SEM images of the untreated external AW showed a significant diversity of the topography in the same image and between the different images. The same diversity was found in the images of the plasma-treated AW. No significant difference was found between the untreated external and the plasma treated AW (data not shown). In conclusion, the SEM analyses emphasized that LPD plasma treatment led to a significant change in the internal surface topography of the AW. It seems that there is an increase in the surface area, which led to increased biofilm attachment. The wettability of AW as well as the bacterial surface hydrophobicity were also examined in order to elucidate whether it is solely the topography that influences the increase in biofilm formation.

**Figure 5:**
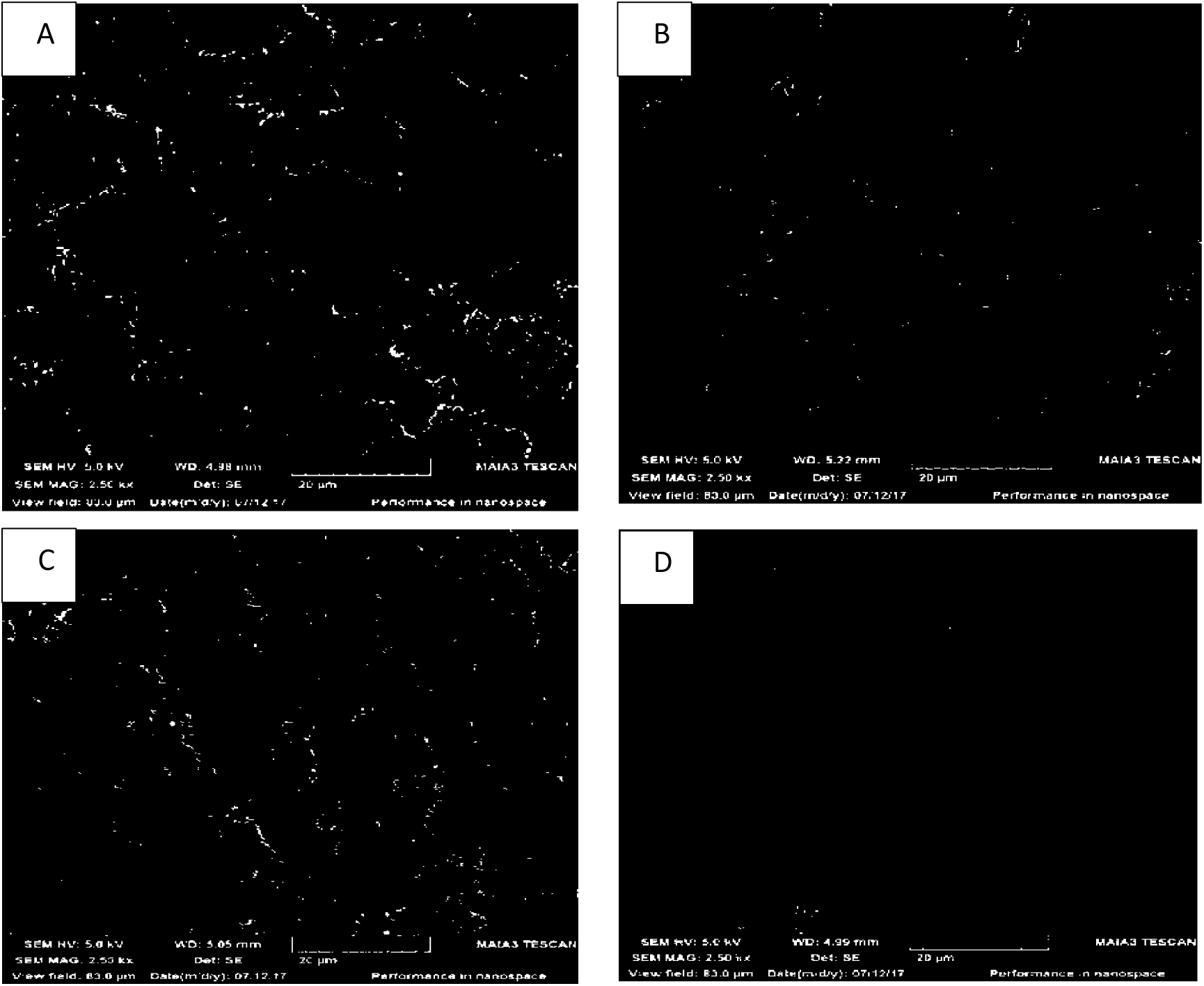
SEM images of untreated (A), low-pressure discharge treated (B), vacuum treated (C) and corona plasma atmospheric discharge treated (D) agricultural waste. The scale bar is 20 µm.

### Wettability of agricultural waste after plasma LPD treatment

Plasma usually activates a surface area and increases its wettability (6). The wettability is characterized by the apparent contact angle of a sessile drop on a surface and the contact angle hysteresis established on the same surface. Surfaces demonstrating obtuse water apparent contact angles are referred as hydrophobic; acute contact angles evidence the hydrophilic nature of a surface (33). The wetting regimes inherent for external and internal surfaces of AW are shown in Figure 6. The apparent water contact angle on untreated internal surface was established as 118°±0.7 (Fig 6 C) and on the untreated external AW it was 75°±2 (Fig 6 D). Note that the internal surfaces demonstrated very high apparent contact angles, close to the maximal theoretically possible ones, inherent for fluorized compounds. Both of external and internal surfaces, that exposed to the LPD were markedly hydrophilized which is expressed by immediately flattered of the sessile drop (Fig 6 E). Contact angle hysteresis established on the external and internal untreated surfaces established with a tilted plate method supplied of value of ca 30°, evidencing more likely the Wenzel-like regime of wetting.

**Figure 6:**
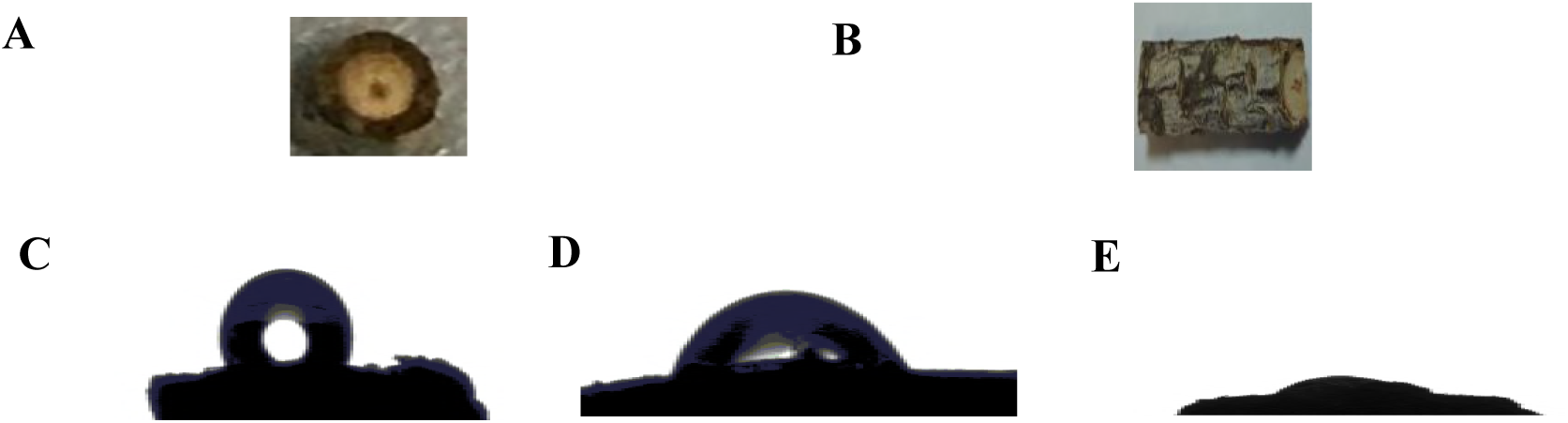
Influence of low pressure discharge on the apparent contact angle of water on agricultural waste is shown. An internal (A) and external agricultural waste surface(B) are depicted. A sessile drop placed on untreated internal (C) and external agricultural waste surfaces is shown (D). (E) A sessile drop placed on the low pressure discharge treated internal surface of agriculture waste is depicted. The complete wetting regimeof wetting is recognized.

### The hydrophobicity of *P. putida* and *B. cereus* cell surface

The bacterial cell surface hydrophobicity can be an important parameter for predicting the adherence of bacteria to a substratum according to its wettability. *P. putida* and *B. cereus* cultures were grown in the presence of toluene as well as glucose and harvested at the end of the log phase. The cultures were washed and the apparent contact angle of a sessile drop on the dried accumulated biomass layer on the filter membrane was measured according to the Van Loosdrecht method (32). The results show very low apparent contact angle values (less than 20°), indicating that the bacterial cell surfaces are hydrophilic (data not shown). Since in this method the filter membrane is blocked very easily we measured the apparent contact angle of water using a very simple procedure. The bacterial cells were washed and prepared according to the Van Loosdrecht method (32) but the washed sediment was resuspended in 200-300 µL of water. The concentrated bacterial suspension was laid on a cover glass and were left to dry for 2h at 30°C. With this method, the apparent contact angle of the sessile drop on the bacterial cell surface was less than 37° (Fig. 7), meaning that the bacterial cell surface of *P. putida* as well as *B. cereus* is hydrophilic.

**Figure 7:**
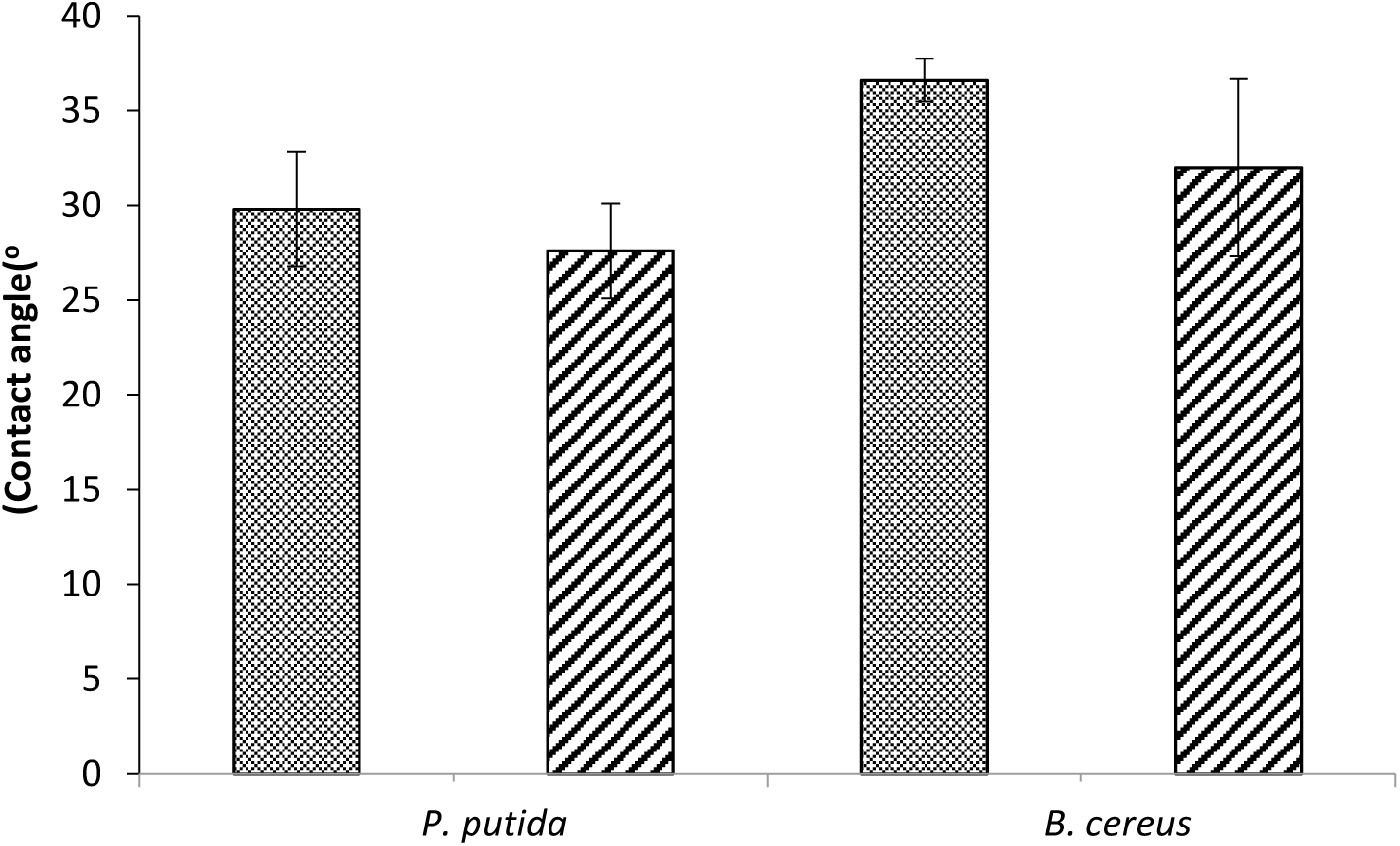
The apparent contact angle of a sessile drop on highly concentrated bacterial ayer placed on a glass slide. When the culture was grown on toluene (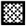) or glucose (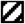).

## Discussion

Biofilm growth may cause a severe problem in industries (34) water systems (35) and hospitals (36). On the other hand, biofilm is useful in a microbial fuel cells for electricity generation [17, 48], microbial electrolysis cells for hydrogen production (38) and in the bioremediation field (20, 30, 39). Bacterial adherence to surfaces is influenced by the bacterial hydrophobicity (32) and by the bacterial physical ability to attach to a substratum using appendages such as the pili, flagella and fimbriae (40). It was reported that surface chemistry, roughness and topography also influence bacterial attachment (23). In recent years there have been many publications reporting different methods of surface modification for enhancing bacterial adherence (25). For example, chamotte porous surface, as well as stainless steel, were modified using organosilane for application in the yeast fermentation industry (41). A carbon nanotubes anode was modified by conducting polymers (Polypyrrole (PPy)-carbon nanotubes (CNT)s and polyaniline (PANI)-CNTs) for application in a microbial fuel cell (42). Carbon nanotubes anode modified using natural-based polymer (chitosan) for application in microbial biosensors technology (43).

In the present study, agricultural waste was treated using LPD and CAPD in an attempt to improve bacterial adherence. The exposure of AW to LPD led to a 3.5-fold increase in biofilm formation of the exogenously added *P. putida* F1 in MMT (minimal medium supplied with toluene) (Fig 2 A) and 1.6-fold in MMG (minimal medium supplied with glucose) compared to the untreated AW (Fig 2 B). When the AW was placed in a sterile MMT or MMG medium, where the biofilm formation evolved from the natural bacteria that attached to the AW, biofilm formation enhancement was 4.2 and 2.7-fold, respectively (Fig 2 A and B). It can be concluded that LPD plasma treatment led to a higher biofilm formation of the exogenously *P. putida* as well as the natural AW bacteria. A similar experiment was performed in the presence of *B. cereus*. Biofilm formation of the exogenously added *B. cereus* in MMT and MMG on LPD-treated AW was 1.3-fold higher than in the untreated AW (Fig 3 A and B). In the sterile MMT and MMG medium, biofilm formation on the LPD-treated AW was 2.8 and 2.4 fold higher, respectively, compared to the untreated AW. The results again show that plasma treatment led to higher biofilm formation.

Surface modification using plasma treatment is considered an economical and environmentally friendly technology, since it is based on mild conditions. The advantages of this technology are that it reduces the degradation of the polymer, changes the surface topography without the use of chemicals, alters the surface physicochemical properties such as hydrophobicity, surface free energy, hydroxyl groups and hydrocarbon content (25). Most of the studies on plasma wood surface modifications are related to the wood industry. Modifying wood surfaces by oxygen plasma at an intensity of 5.8-46.5 kW min m^-2^ treatment was used to accelerate urea formaldehyde, phenol formaldehyde resins and polyurethane coating (44). Wood treatment by air plasma for 1 and 3 s increased the surface wettability and free energy which led to increased adhesion of urea–formaldehyde. It was suggested that the enrichment of the lignin by carboxylic groups contributed to the surface polarity and wettability (45).

In the present research, the plasma wood surface modification was used for the purpose of enhancing biofilm formation for an application in soil bioremediation. As discussed earlier, the plasma LPD treatment led to an increase in biofilm formation of *P. putida, B. cereus* and the natural bacteria. SEM analysis of the plasma LPD-treated AW showed significant topography changes (Fig 5 B) compared to the untreated AW (Fig 5 A). The average diameter of the untreated AW xylem vessels is 10.4±1.4 µm, while in the LPD treated AW it is between 4.4 to 26.4 µm. While in CAPD treated AW the xylem vessels increased to 21.3±4.3 µm and the surface of xylem vessels edge became smooth. It seems that CAPD treatment led to a reduction of the AW surface area. It is important to indicate that while operating the CAPD at a distance of 2 and 5 cm the temperature increased to 107 and 70ºC. We assume that the increase in surface roughness of the LPD-treated AW is one of the reasons for the biofilm enhancement.

Bacterial colonization kinetics on etched surfaces with 10 µm deep and wide of 10 to 40 µm wide pores was dependent on the bacterial strain. However, only motile bacteria were found at the pores’ bottom (23). Nanometric or submicrometric topography were reported to inhibit bacterial adherence, while a micrometric topography increased the available surface for bacterial attachment and biofilm formation (46). Nanopores of 15 and 25 nm on alumina surfaces led to a reduction in bacterial adherence (47). Biofilm formation of the negatively charged surface of *S. aureus* and *P. aeruginosa* was observed on a negatively charged titanium surface. In this case the bacteria were able to overcome the electrostatic repulsion. However, when the titanium surface was changed to a nano-scale topography, a lower biofilm formation was observed (48). No differences in the number of *S. aureus* attachment was found on surface made of polyethylene terephtalate that consisted of nanocylinders of 160 nm height and 110 nm diameter and a smooth surface (49). *P. fluorescens* attachment was higher on a surface with 750 nm wide and 120 nm deep pores, while no colonialization of the cells was observed on the nanometer (amplitude 4-20 nm) structure surface (50).

From our results and from the above-mentioned studies it can be concluded that the surface roughness enhanced biofilm adhesion.

It was reported that the hydrophilicity of the bacteria as well as the substratum surface may also influence bacterial adhesion. The hydrophilicity of *P. putida* as well as *B. cereus* that were grown on toluene or glucose was examined by the apparent contact angle of a sessile drop on a membrane filtrated biomass and was found to be hydrophilic (apparent contact angle of less than 20°). Since the membrane was blocked very easily, we also measured the apparent contact angle of a sessile drop on a layer of concentrated bacterial cells which were laid on a glass slide (Fig 7 The bacterial cell surface was also found to be hydrophilic using this simple method (less than 37°). It was previously reported that the apparent contact angles of water on *E. coli* JM109, D21 and D2 were 19±2, 19±4 and 39±2, respectively. On *B. cepacia* G4, *B. cepacia* Env435 and *B. subtilis* 7003 they were 37±4, 30±3 and 33±2, respectively (51). In contradistinction, the apparent contact angles of water of *P. aeruginosa* and *S. aureus* that were grown in nutrient broth were 43.3±8° and 72.2±8°, respectively (48). The apparent contact angles of water for 23 different bacterial species that were examined were identified as hydrophilic, and no significant differences were found between the hydrophobicity of Gram positive and Gram negative bacteria (32). These data indicate that the bacterial cell surface is mostly hydrophilic, and may also depend on the bacterial medium.

## Conclusions

Biofilm formation of the exogenously added *P. putida* or *B. cereus* on plasma LPD-treated AW was higher than on the untreated AW. We assume that this phenomenon was as a result of the higher roughness as well as the hydrophilic character of the plasma treated AW and the naturally hydrophilic character of the bacterial cell surface.

The low-pressure discharge treatment technology can be further applied for increasing the bioremediation of contaminated soil.

## Acknowledgements

This research was supported in part by the Samaria and Jordan Rift Valley Regional R&D Center and the Research Authority of the Ariel University

